# Episymbiotic bacterium induces intracellular lipid droplet production in its host bacteria

**DOI:** 10.1101/2023.09.06.556576

**Authors:** Pu-Ting Dong, Jing Tian, Koseki J. Kobayashi-Kirschvink, Lujia Cen, Jeffrey S. McLean, Batbileg Bor, Wenyuan Shi, Xuesong He

**Affiliations:** Department of Microbiology, the Forsyth Institute, Boston, MA 02142, USA; Department of Oral Medicine, Infection, and Immunity, Harvard School of Dental Medicine, Boston, MA 02115, USA; Department of Pediatric Dentistry, Peking University School and Hospital of Stomatology, Beijing 100081, China; Broad Institute of MIT and Harvard, Cambridge, MA 02142, USA; Laser Biomedical Research Center, G. R. Harrison Spectroscopy Laboratory, Massachusetts Institute of Technology, Cambridge, MA 02139, USA; Department of Periodontics, University of Washington, Seattle, WA 98195, USA

**Author notes:** Correspondence: Dr. Wenyuan Shi,; Dr. Xuesong He.

## Abstract

*Saccharibacteria* (formerly TM7) *Nanosynbacter lyticus* type strain TM7x exhibits a remarkably compact genome and an extraordinarily small cell size. This obligate epibiotic parasite forms a symbiotic relationship with its bacterial host, *Schaalia odontolytica*, strain XH001 (formerly *Actinomyces odontolyticus* strain XH001). Due to its limited genome size, TM7x possesses restrained metabolic capacities, predominantly living on the surface of its bacterial host to sustain this symbiotic lifestyle. To comprehend this intriguing, yet understudied interspecies interaction, a thorough understanding of the physical interaction between TM7x and XH001 is imperative. In this study, we employed super-resolution fluorescence imaging to investigate the physical association between TM7x and XH001. We found that the binding with TM7x led to a substantial alteration in the membrane fluidity of the host bacterium XH001. Unexpectedly, we revealed the formation of intracellular lipid droplets in XH001 when forming episymbiosis with TM7x, a feature not commonly observed in oral bacteria cells. The TM7x-induced LD accumulation in XH001 was further confirmed by label-free non-invasive Raman spectroscopy, which also unveiled additional phenotypical features when XH001 cells are physically associated with TM7x. Further exploration through culturing host bacterium XH001 alone under various stress conditions showed that LD accumulation was a general response to stress. Intriguingly, a survival assay demonstrated that the presence of LDs likely plays a protective role in XH001, enhancing its overall survival under adverse conditions. In conclusion, our study sheds new light on the intricate interaction between *Saccharibacteria* and its host bacterium, highlighting the potential benefit conferred by TM7x to its host, and further emphasizing the context-dependent nature of symbiotic relationships.

## Introduction

The recently discovered Candidate Phyla Radiation (CPR) contributes to around 26% of bacterial diversity with potentially 73 new phyla (Castelle and Banfield 2018). Among these phyla, *Saccharibacteria* are a group of widespread and genetically diverse ultrasmall bacteria with reduced genomes (McLean et al 2020). Ever since its discovery based on 16S rRNA sequences in 1996 (Rheims et al 1996), *Saccharibacteria* have been detected in a wide range of natural habitats, from the soil, and deep-sea sediments to various human body sites including the gastrointestinal tract, genital tract, and skin (Bik et al 2010, Brinig et al 2003, Fredricks et al 2005, Grice and Segre 2011, Hugenholtz et al 2001, Liu et al 2012). *Saccharibacteria* is particularly prevalent in the oral cavity though with low abundance (Podar et al 2007). However, its relative abundance increases (as high as over 20% of the total oral bacterial community as reported in some studies) in patients with various types of periodontitis (Liu et al 2012, Rylev et al 2011). Research also demonstrated the plausible correlation between *Saccharibacteria* and the occurrence of lung cancer (Lee et al 2016). Like other CPR bacteria, *Saccharibacteria* is known for its recalcitrance to cultivation with limited cultivated strains (McLean et al 2020). As the first cultivated representative of *Saccharibacteria*, “*Nanosynbacter lyticus*” Type-Strain TM7x HMT-952 was isolated from the human oral cavity and enabled the discovery of its parasitic behavior with its bacterial host *Schaalia odontolytica* strain XH001 (He et al 2015, Tian et al 2022).

TM7x is ultrasmall in size (200-300 nm) with a highly reduced genome (705 kb) and lacks amino acid and nucleotide biosynthetic capability (He et al 2015). It is an obligate epibiont parasite living on the surface of and inducing stress in its bacterial host cell XH001 (Bor et al 2016, Utter et al 2020). A recent study demonstrated that through a complete arginine deiminase system, an arginine catabolism pathway acquired during its environment-to-mammal niche transition, TM7x can catabolize arginine, produce adenosine triphosphate and ammonia, then confers a beneficial effect towards its host cells XH001, especially in the low-pH environment (Tian et al 2022). A comprehensive transcriptome study from TM7x and XH001 provides mechanistic insights into this episymbiotic lifestyle in the level of gene expression (Hendrickson et al 2022).

In this study, we utilized super-resolution fluorescence microscopy to offer nanoscale insights into the physical interactions between TM7x and XH001. Notably, the infection of TM7x had a significant impact on the membrane fluidity of host cells XH001, resulting in a distinct distribution of liquid-ordered and disordered phases compared to the organized distribution observed in XH001 cells alone. Additionally, we made an intriguing discovery of the enhanced production of intracellular lipid droplets (LDs) in the XH001 cells when physically associated with TM7x. Furthermore, we employed label-free non-invasive Raman spectroscopy to capture multiple phenotypic differences based on the Raman spectra, demonstrating saturated fatty acids being the most prominent contributor to these signature differences. Intriguingly, XH001 cells alone also exhibited the accumulation of saturated fatty acids when subjected to various stress conditions, indicating that LDs formation could be a general response to stress and serves as a stress marker. The subsequent starvation assay further demonstrated that the accumulation of fatty acids likely plays a beneficial role in protecting XH001 cells against stress factors and enhancing their survival. Overall, the combination of advanced imaging techniques and label-free Raman spectroscopy revealed some of the emergent features in XH001 when forming symbiosis with TM7x and provided valuable new insights into this intriguing bacterial interspecies interaction.

## Results

### Super-resolution fluorescence imaging of BODIPY C_1_, C_12_-labeled XH001 and XH001/TM7x

Previous studies have shown that TM7x predominantly resides on the surface of its host bacterium XH001 through fluorescence in-situ hybridization (FISH) imaging (Bor et al 2016, He et al 2015) and phase contrast microscopy. However, the morphology of TM7x remains difficult to resolve as the size of the TM7x cells is around 200-300 nm, close to the diffraction limit (Rust et al 2006). Microscopic approaches with sub-diffraction-limit capacity are needed. Through super-resolution fluorescence imaging achieved by confocal fluorescence microscopy (**Materials and Methods**) with an airyscan detector (Huff 2015), a lateral resolution of 120 nm can be achieved (Wu and Hammer 2021). To visualize both TM7x and host cells XH001, we utilized a fluorescence dye, BODIPY C_1_, C_12_ (**Materials and Methods**) which labels every organelle that is hydrophobic (Kaiser and London 1998). We compared the image quality of BODIPY C_1_, C_12_-stained XH001/TM7x under the conventional confocal microscope and confocal microscope with airyscan detector, respectively. As shown in **Figure 1A**, fluorescence images under a confocal microscope with airyscan detector apparently exhibit higher image resolution. Moreover, BODIPY can label both XH001 cells and attached TM7x cells.

**Figure 1.**
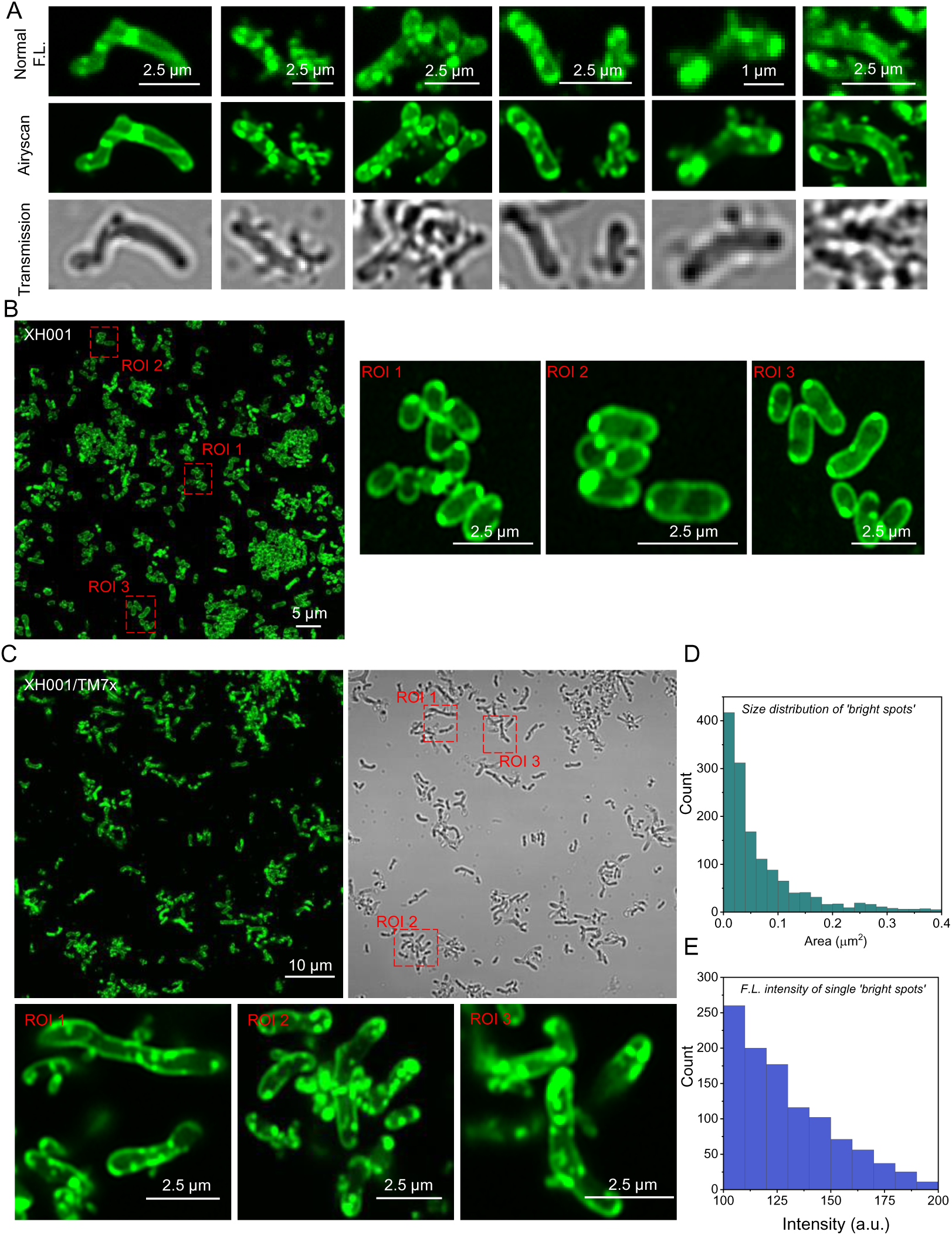
Confocal laser scanning imaging of BODIPY C_1_, C_12_-labeled monoculture XH001 cells and co-culture XH001/TM7x with airyscan detector. **A**. Comparison of the fluorescence images acquired under the conventional confocal microscope and confocal microscope with airyscan detector. **B**-**C**. Super-resolution fluorescence imaging of host cells XH001 and XH001/TM7x along with zoom-in views of three regions of interest (ROIs), respectively. **D**. Size distribution of ‘bright dots’ from **C**. **E**. Fluorescence intensity distribution of individual ‘bright spots’ from **C**. Fluorescence images were obtained after labeling XH001 or XH001/TM7x with BODIPY for 30 min and sandwiched between a cover glass and cover slide. Scalar bars are labeled directly on the corresponding images.

We then compared the morphology of BODIPY C_1_, C_12_-labeled monoculture XH001 cells, and XH001/TM7x co-culture under the confocal microscope with airyscan detector. Monoculture XH001 cells displayed fluorescence signals mainly from the cell membrane and some from bright dots that were mostly associated with the membrane (**Figure 1B**), presenting a rod-shaped morphology. Intriguingly, in the XH001/TM7x co-culture, XH001 cell morphology is highly irregular, and fluorescence signals are from both the cell membrane and mostly intracellular ‘bright dots’ (**Figure 1C**). These ‘bright dots’ showed an average area of 0.099 µm^2^, corresponding to a 340-nm diameter dot size (**Figure 1D**). And they demonstrated excellent fluorescence intensity (**Figure 1E**) compared to the fluorescence stain from the cell membrane. It was also known that BODIPY C_1_, C_12_ has been widely employed to visualize neutral lipids or lipid droplets (Qiu and Simon 2016, Rumin et al 2015). We thus suspected that these intracellular ‘bright dots’ might be lipid droplets (LDs).

### Laurdan fluorescence imaging suggests the host cell membrane fluidity change induced by TM7x

In addition to cell morphology change, one of the characteristic features of episymbiosis is the intimate physical interaction between epibiont and its host cell. To further understand the cell membrane physiological change in host bacterium XH001 induced by TM7x, Laurdan was employed to characterize the membrane fluidity and organization. Laurdan, a membrane fluorophore sensitive to the local membrane packing, has been extensively utilized to quantify the degree of lipid packing and membrane fluidity due to the dipolar relaxation effect (Sanchez et al 2012, Weber and Farris 1979). Under an excitation wavelength of 405 nm, the emission spectrum of Laurdan peaked at 490 nm when the membrane lipids are in a disordered phase (liquid disordered phase, more fluid) and shifted to ∼ 440 nm when the membrane lipids are in a more packed situation (liquid-ordered phase, more rigid) (Golfetto et al 2013, Gunther et al 2021, Kaiser et al 2009). Then a generalized polarization (GP) value can be derived to quantify the membrane fluidity:

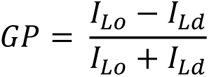

where 𝐼_𝐿𝑜_, 𝐼_𝐿𝑑_ is the fluorescence intensity from liquid ordered phase and liquid disordered phase, respectively. Therefore, the obtained GP values range from -1 (being least ordered, very fluid) to +1 (being most ordered, very rigid).

To image and quantify membrane fluidity at the nanometer scale, super-resolution fluorescence imaging with an airyscan detector was employed to harvest the fluorescence images from liquid disordered phase and liquid-ordered phase by choosing corresponding emission filters (**Materials and Methods**). As shown in **Figure 2A**, the liquid-ordered phase fluorescence signal of XH001 cells is mainly from the parallel edges, whereas liquid-disordered phase fluorescence is mostly from the distal ends of cells. In contrast, when forming symbiosis with TM7x, XH001 cells displayed more randomly distributed fluorescence signals from two channels, and the membrane patches of liquid-ordered/disordered phase seem to be stochastically allocated (**Figure 2B**).

**Figure 2.**
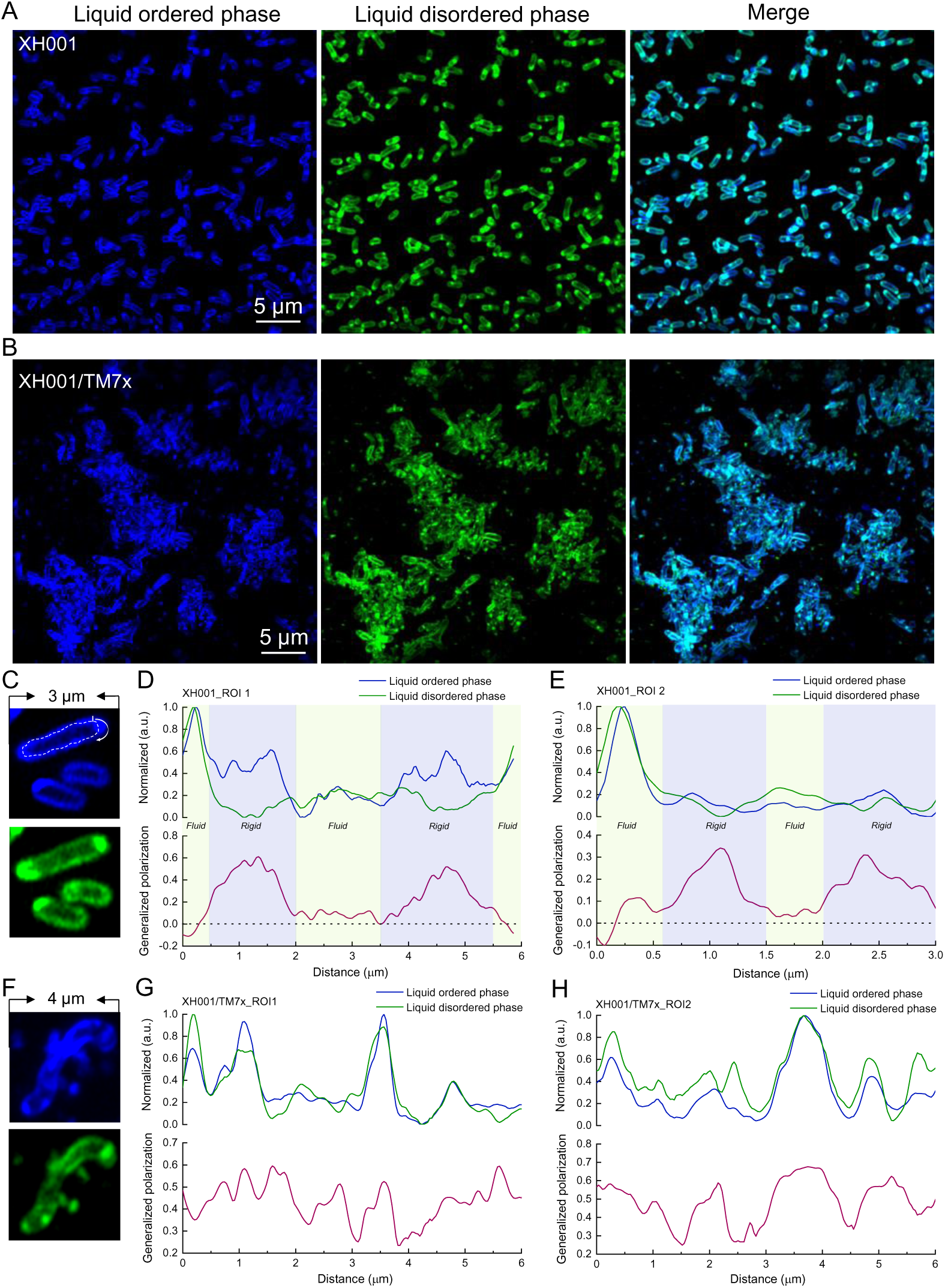
Mapping membrane fluidity of XH001 and XH001/TM7x through super-resolution fluorescence imaging of Laurdan-labeled XH001 and XH001/TM7x. **A-B**. Super-resolution fluorescence images of liquid ordered phases and liquid disordered phases from XH001/TM7x and XH001, respectively. Excitation laser: 405 nm. Fluorescence signal from liquid ordered/disordered phase was collected by an optical bandpass filter (410-460 nm) and (470 nm–550 nm), respectively. Scalar bar = 5 µm. **C-E**: Zoom-in view of Laurdan-labeled XH001 cells along with single-cell mapping of generalized polarization (GP). **F**-**H**: Zoom-in view of Laurdan-labeled XH001/TM7x along with single-cell mapping of GP. A GP value of 0 was highlighted by black dashed lines.

GP values of XH001 cells from monoculture and XH001/TM7x co-culture were then obtained based on the above equation. Here, to better understand the distribution of lipid patches from liquid-ordered and disordered phases along the cell membrane, circular GP values of the host cell membrane were calculated starting from the distal end of the cell (**Figure 2C**). As shown in **Figure 2D**-**E**, the central parts of XH001 cells are more rigid with a GP value around 0.3-0.6. The distal ends of host cells exhibit smaller GP values (0-0.1), indicative of fluid lipids clustered together at the ends of XH001 cells. This evidence suggests that lipid packing in XH001 cells from monoculture is highly oriented. Interestingly, XH001 cells forming symbiosis with TM7x displayed a highly randomized lipid packing (**Figure 2, F-H**) as GP value has no specific oscillation pattern along the cell peripheral. This data suggests that symbiosis with TM7x drastically affects the organization of lipid molecules in XH001 cell membrane. In short, through Laurdan fluorescence imaging, we showed that membrane fluidity of host cells XH001 is significantly altered as a result of TM7x infection: once ordered lipid packing becomes highly randomized. Meanwhile, as a hydrophobic dye, Laurdan staining also revealed intracellular LD-like structures in XH001 when forming symbiosis with TM7x.

### LipidSpot staining consolidates the formation of intracellular lipid droplets (LDs) in XH001 when forming symbiosis with TM7x

Lipid droplet (LD), a monolayer phospholipid membrane-bound organelle found in almost all eukaryotes, plays essential roles in cellular lipid homeostasis (Gao and Goodman 2015). However, bacterial intracellular lipid droplets (LDs) are less common and remain underexplored. To confirm these BODIPY and Laurdan-stained ‘bright spots’ are indeed LDs, we utilized an LDs-specific dye, ‘LipidSpot (Farmer et al 2020)’, to stain XH001 and XH001/TM7x (**Materials and Methods**). Confirming our previous staining, we observed lots of lipid droplets showing up in the co-culture XH001/TM7x from different fields of view (**Figure 3A**). On the contrary, very faint signals can be detected in XH001 cells from monoculture (**Figure 3B**), especially under different color bars. A higher magnification view further consolidates this phenomenon (**Figure 3, C-D**). Intriguingly, all of these ‘lipid droplets’ are localized intracellularly, at the cell poles or associated with cell membrane. And TM7x alone was not labeled by this dye. To investigate the spatial arrangement of LDs in relationship to TM7x attachment, we also counterstained TM7x using fluorescence in situ hybridization in the lipidspot-stained co-culture XH001/TM7x. As shown in **Supplementary Figure 1**, TM7x closely correlates with LDs spatially (with a pair correlation value larger than 1), which suggests LD tends to form in the area that is spatially close to the associated TM7x cells. Collectively, data from BODIPY- and Laurdan-stained images as well as LipidSpot-labeled fluorescence images strongly indicated that the episymbiotic interaction with TM7x significantly induces the production of intracellular LDs.

**Figure 3.**
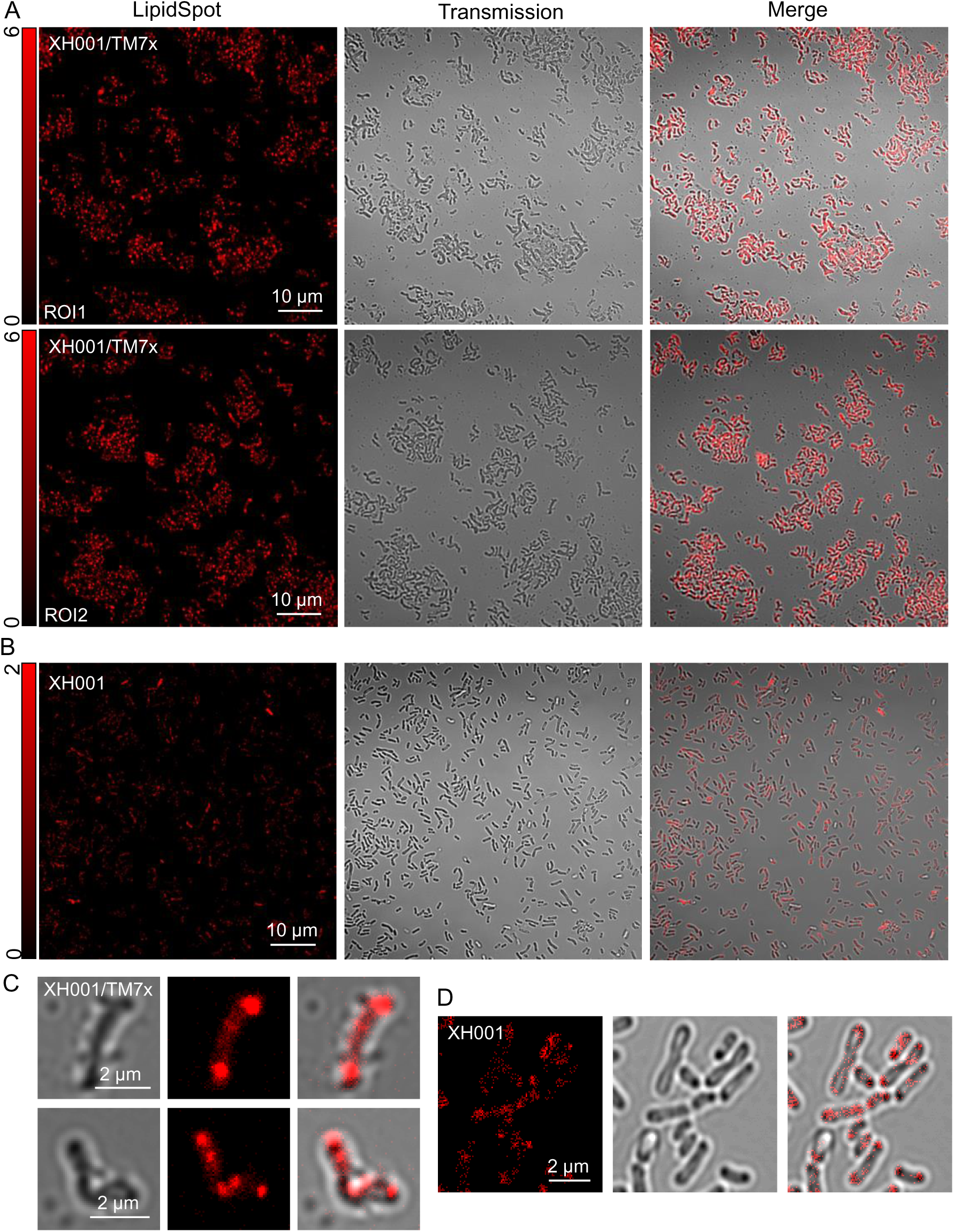
Confocal laser scanning imaging of LipidSpot-labeled XH001/TM7x and XH001. **A-B**. Fluorescence images, transmission images, and merged images of LipidSpot-labeled XH001/TM7x from two ROIs and XH001 cells, respectively. Scalar bar = 10 µm. **C**-**D**. Zoom-in view of LipidSpot-labeled XH001/TM7x and XH001 cells, respectively.

### Raman spectroscopy reveals the enhanced production of LDs in XH001 when associated with TM7x

BODIPY C_1_, C_12_, Laurdan, and LD-specific fluorescence imaging collectively indicated an increase in LD production within XH001 cells when associated with TM7x. To further validate these observations, we sought to explore alternative methodologies, such as Raman spectroscopy, to acquire independent confirmation. Moreover, this non-invasive assay provides a comprehensive perspective on the shift in cellular metabolites within XH001 cells upon interaction with TM7x.

Raman spectroscopy has the capability to probe the chemical compositions of biological analytes in a label-free and non-invasive manner (Cheng and Xie 2015, Du et al 2020, Wachsmann-Hogiu et al 2009, Xu et al 2019). Raman scattering is a process of inelastic scattering of photons by molecular bonds given that every chemical bond inside a specific molecule has distinctive vibrational energy. We thus wondered how different these intracellular biomolecules are inside XH001 cells when being TM7x-free vs. attached to TM7x. To answer this question, cells from XH001 monoculture and XH001/TM7x co-culture were fixed inside 10% formalin, washed with milli-Q water, and then air-dried onto an aluminum substrate. Raman spectra of XH001 cells and XH001/TM7x were then acquired under a HORIBA Raman spectrometer (**Materials and Methods**).

As depicted in **Figure 4A** and **Supplementary Figure 2**, the fingerprint region spanning from 320 cm^-1^ to 1800 cm^-1^ effectively captures the majority of vibrational data related to intracellular biomolecules, revealing the distinctive and observable Raman phenotype characteristic of cells. Following vector normalization (**Materials and Methods**), the Raman spectra of both XH001 and XH001/TM7x cells yielded the identification of several typical Raman peaks: 720 cm^-1^, 780 cm^-1^ (DNA or RNA), 745 cm^-1^ (cytochrome c), 1003 cm^-1^ (phenylalanine), 1128 cm^-1^ (saturated lipids) (Czamara et al 2015), 1660 cm^-1^/1250 cm^-1^ (Amide I/III, protein). Notably, the Raman intensities of certain peaks, such as cytochrome c, exhibited significant disparities. Importantly, these variations cannot be solely attributed to the additive presence of TM7x (**Supplementary Figure 2**).

**Figure 4.**
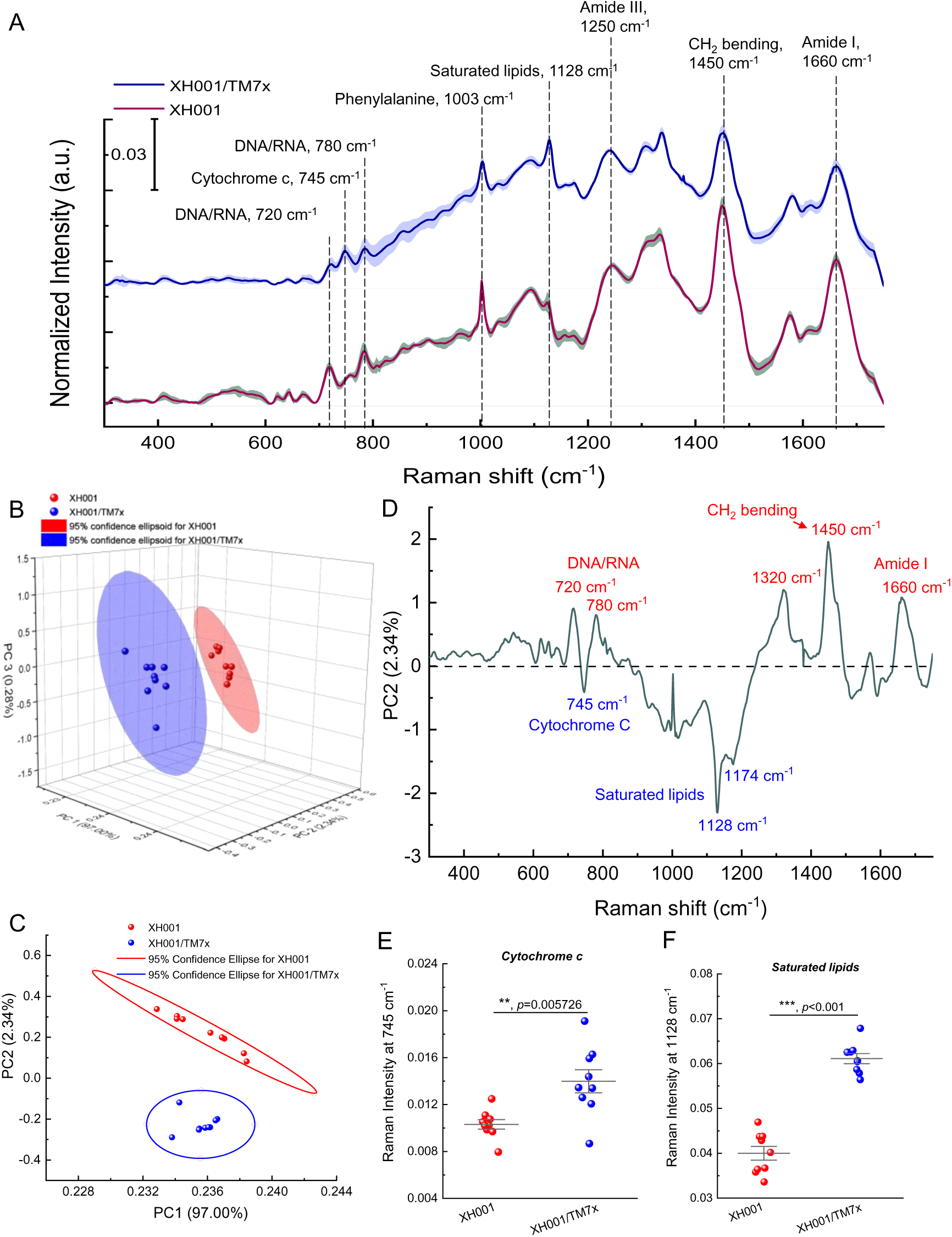
Raman spectroscopy of XH001 and XH001/TM7x along with the corresponding principal component analysis. **A**. Raman spectra of XH001 cells and XH001/TM7x from three biological replicates after vector normalization. Data: Mean ± Standard deviation (SD). Interested Raman peaks were highlighted by gray dashed lines. **B**. Three-dimensional principal component analysis (PCA) plot of Raman spectra of XH001 cells and XH001/TM7x. Each dot represents a Raman spectrum. Data are from three biological replicates. **C**. Two-dimensional principal component analysis plot from principal component 1 (97.00% of the total variation) and 2 (2.34% of the total variation). 95% confidence intervals were represented by ellipses. **D**. Spectral information of PC2. Biomolecules with enhanced production in the case of XH001/TM7x were highlighted in blue and XH001 cells in red. **E**-**F**. Quantitative and statistical analyses of Raman intensities from cytochrome c (750 cm^-1^) and saturated lipids (1128 cm^-1^). Statistical analysis was conducted through a two-tailed student unpaired *t*-test. ***: *p*<0.001.

To provide a comprehensive representation of the Raman spectral distinctions, we employed principal component analysis (PCA). PCA, a powerful statistical technique, enables the reduction of data dimensions while retaining the majority of the variance present in the original dataset (Jolliffe and Cadima 2016). Subsequently, we generated two-dimensional and three-dimensional plots, utilizing different combinations of scores from the first two or three components, respectively. In the context of the three-dimensional plot (**Figure 4B**), a divergence emerges between the Raman spectra of XH001 in monoculture and those of XH001/TM7x. Notably, the two-dimensional PCA plot (**Figure 4C**), based on the first two PCs, demonstrates a substantial and statistically significant disparity in the PC2 scores between XH001 and XH001/TM7x (*p*<0.001, student unpaired *t*-test).

To decipher which specific biomolecules within PC2 contribute significantly to the discernible distinctions in Raman spectra between XH001 monoculture and XH001/TM7x co-culture, we extracted the spectral information associated with PC2, responsible for elucidating 2.34% of the total variation. As depicted in **Figure 4D**, Raman peaks at 745 cm^-1^, 1128 cm^-1^, and 1174 cm^-1^ stand out in the case of XH001/TM7x in comparison with XH001. These Raman peaks correspond to cytochrome c (745 cm^-1^) and saturated lipids (1128 cm^-1^ and 1174 cm^-1^) (Baron et al 2020, Czamara et al 2015), respectively. Furthermore, the spectral details of PC2 unveil additional significant disparities encompassing nucleic acids (720 cm^-1^/780 cm^-1^) and amide I (1660 cm^-1^).

There was a difference not only in the pattern of the peaks but also in the intensity of the peaks. Therefore, we performed a quantification of intracellular biomolecules by integrating Raman bands at 745 cm^-1^ (cytochrome c, **Figure 4E**) and 1128 cm^-1^ (saturated fatty acids, **Figure 4F**). This quantification approach is underpinned by the linear relationship between Raman intensity and the concentration of biomolecules (Jones et al 2019). We found that there is a significantly higher signal (*p*<0.001) in the Raman spectra of XH001/TM7x compared to XH001 regarding the content of cytochrome c and saturated lipids. In particular, the augmented production of saturated fatty acids, which are primary constituents of LDs (Wältermann and Steinbüchel 2005), from XH001/TM7x co-culture is consistent with our abovementioned microscopic evidence.

### XH001 cells exhibit enhanced accumulation of LDs in the presence of various stress factors

According to a previous report (Hendrickson et al 2022), TM7x induces stress in XH001 cells, particularly in the stable symbiosis state. Furthermore, it has been demonstrated that prokaryotes possess the capacity to accumulate lipophilic compounds within the cytoplasm, forming lipid inclusions (Wältermann and Steinbüchel 2005). These lipophilic compounds, such as polyhydroxybutyrate, triacylglycerol, and wax ester, are accumulated in response to stress (e.g., low nitrogen availability) (Zhang et al 2022). We wondered whether the formation of LDs observed through Raman spectroscopy and fluorescence microscopy was due to stress-induced host response. To explore this phenomenon, we cultured XH001 cells under diverse stressed conditions, including distinct oxygen levels (0% and 21%), which have been documented to elicit varying degrees of stress responses in XH001 cells (Bor et al 2016). We also subjected XH001 cells to treatments with various H_2_O_2_ concentrations.

As illustrated in **Figure 5A**, the minimum inhibitory concentration (MIC) of H_2_O_2_ against XH001 cells is 44 µM. To maintain viability while inducing stress, we opted for a sub-MIC concentration of 22 µM to treat XH001 cells. In all, XH001 cells were treated with 22-µM H_2_O_2_, or grew under 0% O_2_, or 21% O_2_, and subsequently fixed in 10% formalin and analyzed by Raman spectra to assess potential phenotypic differences. Notably, distinct Raman peaks at 745 cm^-1^ (cytochrome c), 1128 cm^-1^ (saturated fatty acids), and 1660 cm^-1^ (Amide I, protein) were evident in the groups subjected to different stress (**Figure 5B**) compared to XH001 cells cultured under the optimized condition (2% O_2_). Global differences between XH001 cells cultured under optimized and stressed conditions were revealed through two-dimensional principal component analysis (**Figure 5C**), with principal component 2 illustrating the most significant difference and all stress conditions clustered together with XH001/TM7x co-culture compared to XH001 alone (**Supplementary Figure 3**). Spearman’s correlation analysis also demonstrated that saturated lipids display a significant positive correlation with cytochrome c (**Figure 5D**), and a negative correlation with Amide I (**Figure 5E**). This likely indicates that the increased amount of cytochrome c may boost the accumulation of saturated fatty acids at the expense of cellular amino acids or protein. Additionally, we examined the formation of lipid droplets using super-resolution fluorescence microscopy. **Figure 5F** displays BODIPY C_1,_ C_12_-labeled XH001 cells under the anaerobic cultured condition and revealed apparent lipid droplet accumulation, especially when comparing XH001 cells cultured under the microaerophilic condition (**Figure 1B**).

**Figure 5.**
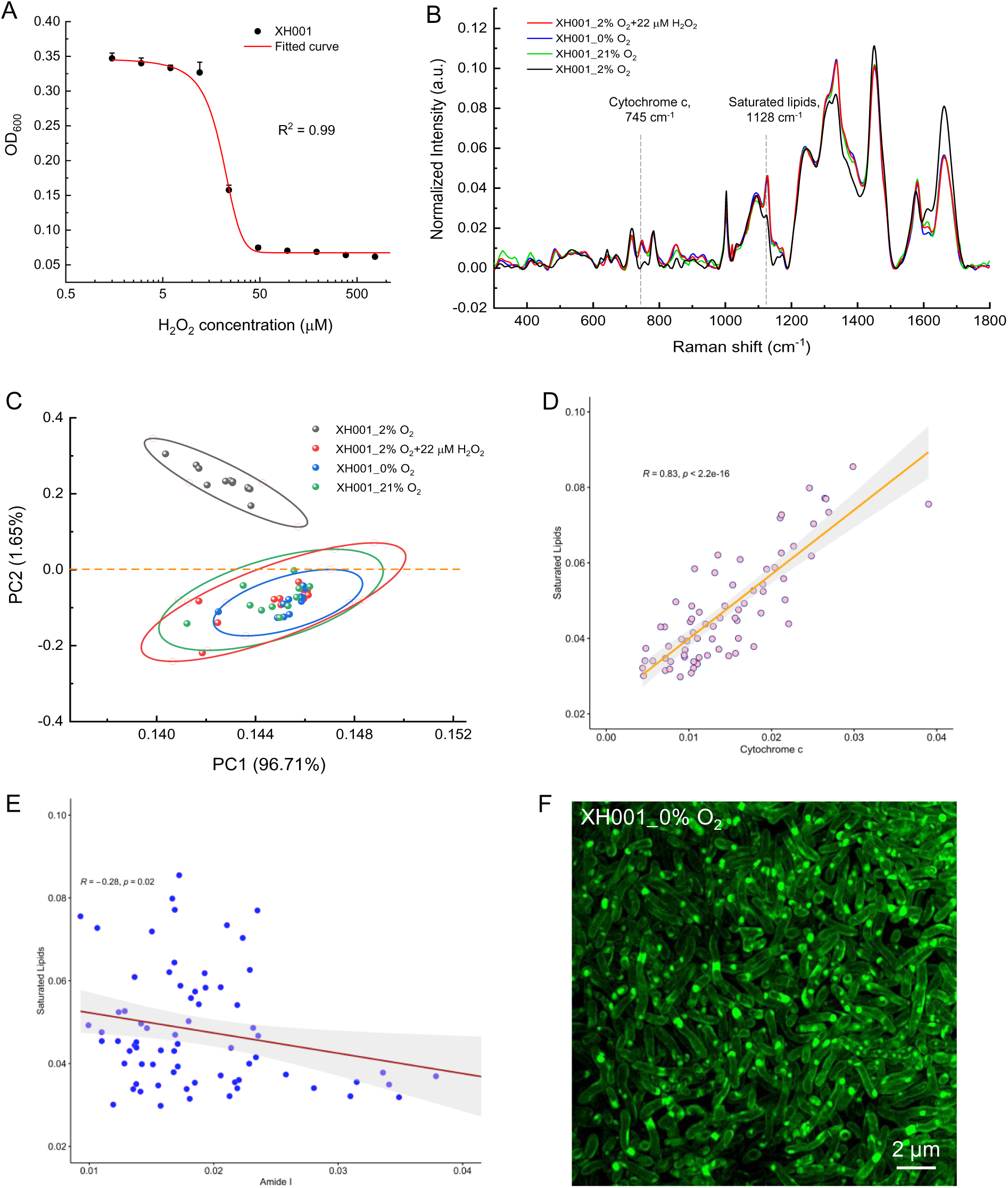
Characterization of XH001 cells under various stressed conditions by Raman spectroscopy and super-resolution fluorescence imaging. **A.** Minimum inhibitory concentration (MIC) of H_2_O_2_ against host cells XH001 alone. Data was fitted by a dose-response relationship function. **B**. Raman spectra of XH001 cells cultured under various stressed conditions, with each spectrum averaged from at least ten individual plots of three biological replicates. **C**. Two-dimensional principal component analysis plot of Raman spectra acquired from XH001 cells under different stressed conditions. Plots were depicted from principal component 1 (96.71% of the total variation) and 2 (1.65% of the total variation). 95% confidence intervals were represented by ellipses. **D**. Correlation analysis between cytochrome c (745 cm^-1^) and saturated lipids (1128 cm^-1^). Spearman’s correlation coefficient and *p*-value were labeled in the plots. **E**. Correlation analysis between Amide I (1660 cm^-1^) and saturated lipids (1128 cm^-1^). Spearman’s correlation coefficient and *p*-value were labeled in the plots. **F**. Super-resolution fluorescence imaging of BODIPY C_1,_ C_12_-labeled XH001 cells cultured inside the anaerobic conditions. Scalar bar = 2 µm.

### Accumulation of LDs enhances cell survival under stressed conditions

To investigate the biological implications of increased accumulation of fatty acids, we conducted a starvation assay as follows (**Materials and Methods**): XH001 cells cultured under both optimized (XH001_normal) and pre-stressed (0% O_2_, overnight) conditions (XH001_prestress) were extensively washed to remove residual medium and resuspended in 1×PBS and incubated in the microaerophilic chamber (2% O_2_). The temporal changes in colony-forming unit (CFU) assay, Raman spectra, and H_2_O_2_ susceptibility test were determined for each culture. The first drastic difference was observed in CFU where normal and prestressed XH001 groups began to display separation at 20 hours post starvation (**Figure 6, A-B**). XH001_normal group suffered significantly more (three orders of magnitude) reduction in CFU compared to the pre-stress group at 48 hours post-starvation, suggesting that pre-stress enhances the survival of XH001 cells under starvation conditions. Moreover, the H_2_O_2_ susceptibility test between the two groups provided further evidence that pre-stress significantly boosts XH001 cells’ resilience against oxidative environments (**Figure 6C**). Intriguingly, Raman spectroscopy analysis demonstrated that the amount of saturated lipids decreases as the starvation process endures in the group XH001_prestress (**Figure 6, D-E**), and this is positively correlated with the time-course decrease in CFU of pre-stressed XH001 cells, which further consolidates the protective role of saturated fatty acids. We also noted the change in the intracellular amount of other biomolecules (**Supplementary Figure 4**), such as amino acids (1660 cm^-1^) and nucleic acids (DNA or RNA, 780 cm^-1^), which suggests fatty acids accumulation may not be the only factor contributing to the cell survival under stressed conditions.

**Figure 6.**
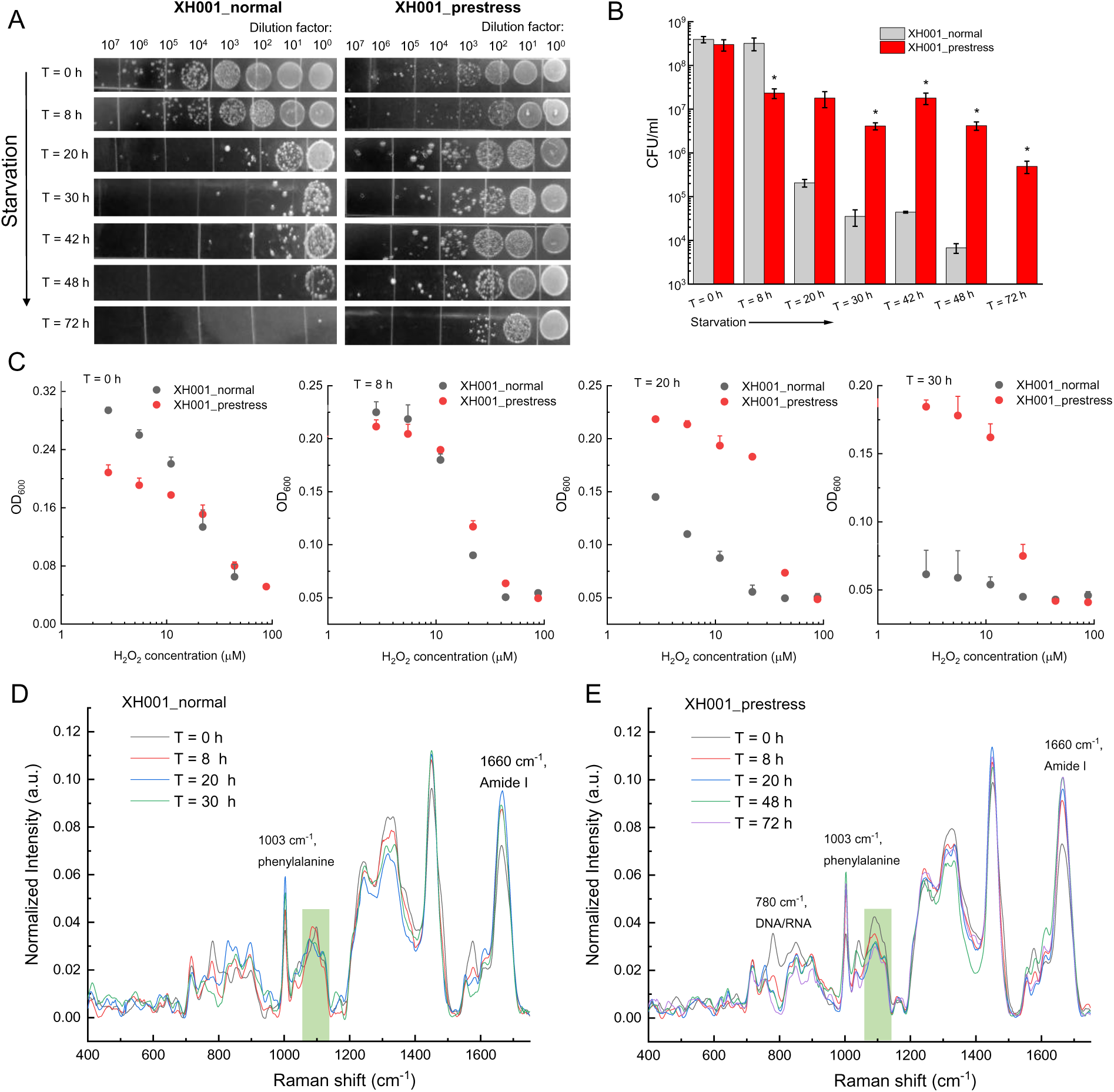
Saturated fatty acids potentially contribute to the survival of XH001 cells under stressed conditions. **A**. Colony-forming units (CFU) of XH001 cells with and without pre-stress under time-course starvation conditions. **B**. Quantification analysis of CFU from **A**. Data: Mean ± Standard deviation from three replicates. Statistical analysis was conducted between the normal- and pre-stressed groups by a student unpaired *t*-test. *: *p* < 0.05. **C**. Dose-response of H_2_O_2_ against XH001 cells under both normal and prestressed conditions at different starvation times. Mean + Standard deviation from three biological replicates. **D-E**. Raman spectra from XH001 cells under both normal (**D**) and prestressed conditions (**E**). Data were averaged from at least ten spectra acquired from three biological replicates. Regions of interest were highlighted by green rectangular boxes.

### Other bacteria do not exhibit an apparent accumulation of LDs under stressed conditions

Next, we wondered whether the accumulation of saturated fatty acids and the formation of intracellular LDs when under stress is a general feature of human-associated bacteria. We examined the characteristics of other bacteria, such as *Fusobacterium nucleatum* (*Fn*), *Streptococcus mutans* (*S. mutans* UA159), as well as a pathogenic *Staphylococcus aureus* strain, MRSA USA300, in the presence of sub-MIC H_2_O_2_ by Raman spectroscopy and super-resolution fluorescence microscopy.

Firstly, we assessed the H_2_O_2_ susceptibility of *Fn* ATCC 10953 (*Fn* 10953), *Fn* ATCC 23726 (*Fn* 23726), *S. mutans*, and MRSA USA300 (**Figure 7, A-D**). Subsequently, we selected a sub-MIC concentration of H_2_O_2_ to induce stress in the four bacteria. Raman spectra were then acquired from fixed bacteria, both with and without the addition of H_2_O_2_. Unlike XH001 cells under stress, those bacteria do not exhibit increased Raman intensity at 1128 cm^-1^, indicating a lack of accumulation of saturated fatty acids (**Figure 7, E-H**).

**Figure 7.**
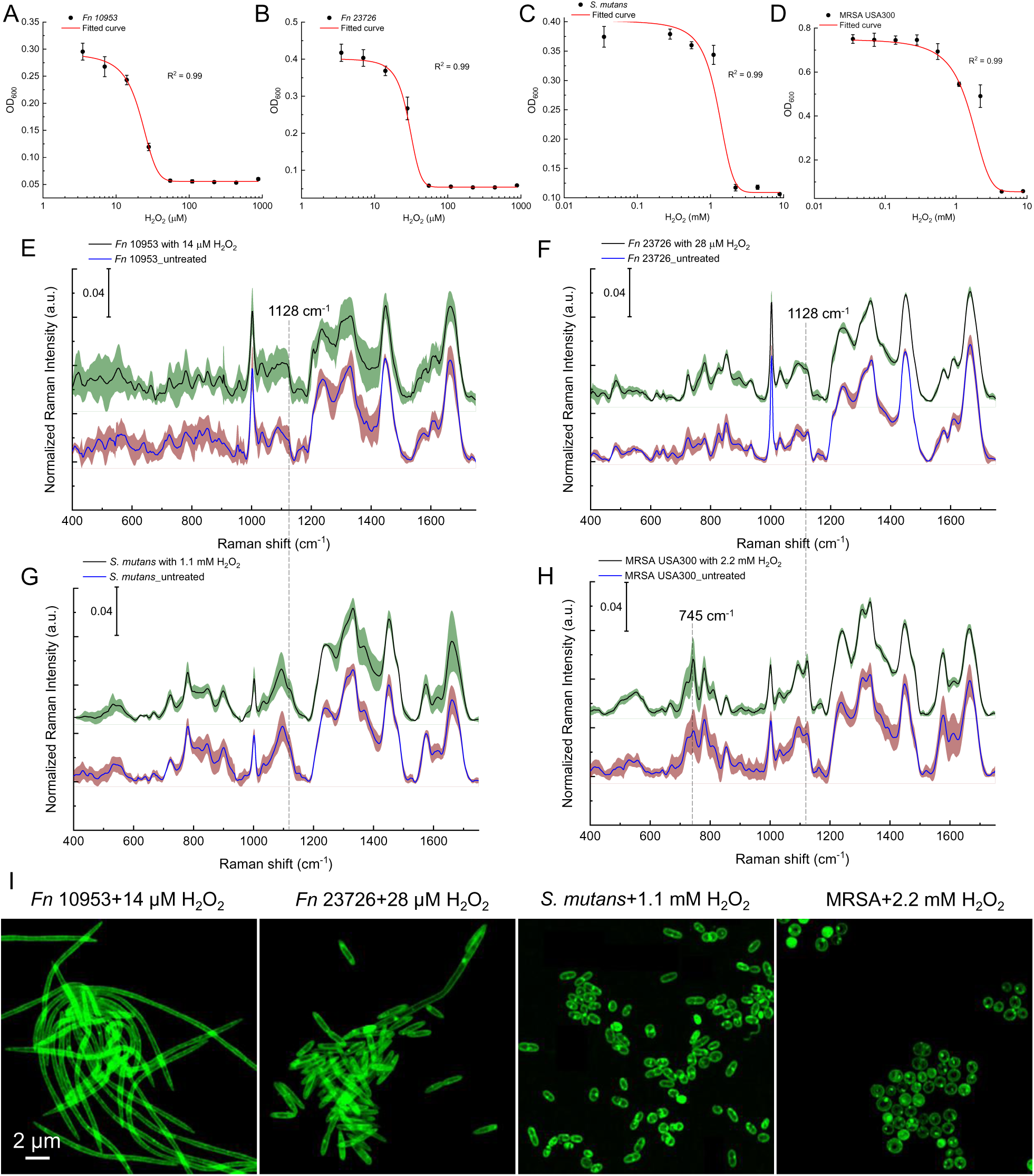
Characterization of other bacteria under stressed conditions by Raman spectroscopy and super-resolution fluorescence microscopy. **A-D:** Dose-response of H_2_O_2_ against *Fn* 10953, *Fn* 23726, *S. mutans*, and MRSA USA300. **E-H**: Raman spectra of *Fn* 10953 (**E**), *Fn* 23726 (**F**), *S. mutans* (**G**), and MRSA USA300 (**H**) with and without the treatment of H_2_O_2_. Data: Mean ± Standard deviation (shadow) from three biological replicates. Raman peaks at 745 cm^-1^ and 1128 cm^-1^ were highlighted by dashed lines. **I**: Super-resolution fluorescence imaging of BODIPY C_1_, C_12_-labeled *Fn* 10953, *Fn* 23726, *S. mutans*, and MRSA after the treatment of sub-MIC H_2_O_2_. Scalar bar = 2 µm.

In the case of MRSA USA300 with and without the treatment of sub-MIC H_2_O_2_, we noticed an increase with regards to Raman intensity at 745 cm^-1^ and 1128 cm^-1^ (**Figure 7H**). It has been reported that both saturated lipids and cytochrome c also contribute to the Raman peak at 1128 cm^-1^ (Baron et al 2020, Pezzotti et al 2022, Xu et al 2023). To discern the impact of cytochrome c upon the Raman intensity at 1128 cm^-1^, we then quantified the integral Raman intensity of cytochrome c and saturated lipids from both XH001 and MRSA cells under stressed conditions. As illustrated in **Supplementary Figure 5**, a notable three-fold difference emerged between cytochrome c and saturated lipids in the context of XH001 cells subjected to stressful conditions (*p*<0.0001). In contrast, no significant distinction was discerned for MRSA under stress conditions (*p*=0.06). This observation strongly suggested that, in the case of MRSA, the Raman intensity at 1128 cm^-1^ predominantly originated from cytochrome c, as indicated by the pronounced Raman intensity at 745 cm^-1^. Moreover, a subsequent super-resolution BODIPY C_1_, C_12_ imaging analysis revealed the absence of discernible LDs formation in these bacteria when exposed to stressful conditions. Hence, this compelling evidence hints at the notion that the accumulation of saturated fatty acids may represent a unique hallmark of *Actinomyces*.

## Discussion

Illustrating the physical interaction between *Saccharibacteria* and its host bacteria is essential to understanding this intimate, yet understudied interspecies interaction. However, current microscopic images have limited resolution to clearly depict this episymbiotic relationship. Here, through super-resolution fluorescence imaging, the morphology of co-culture XH001/TM7x was clearly discernible with around 120-nm lateral resolution under BODIPY staining (**Figure 1**). Rod-shaped host cells XH001 become irregular shapes in the co-culture XH001/TM7x, which might be related to the lower expression of chromosomal partitioning genes (Hendrickson et al 2022). Laurdan quantification indicated that the cell membrane fluidity or lipid packing of host cells XH001 was drastically affected due to the association with TM7x: from highly ordered to randomly distributed (**Figure 2**). We discovered the formation of intracellular LDs in XH001 cells in the co-culture (**Figure 1 and 2**). LipidSpot assay and label-free Raman spectroscopy assay further consolidated this discovery (**Figure 3 and 4**). The data implies that epibiont *Saccharibacteria* significantly alters the host cell membrane organization and elicits the cellular metabolic response, leading to the accumulation of saturated fatty acids and the enhanced formation of LDs.

LDs are a unique multi-functional organelle comprising a phospholipid monolayer surrounding a hydrophobic neutral lipid core that contains cholesterol esters, triacylglycerols, and wax esters (37). They are ubiquitous lipid storage organelles existing in all eukaryotic cells (38, 39), especially when the cells are under stress or challenge. Intracellular LDs are less common in bacteria, with limited reports. Gram-positive actinobacteria and Cyanobacteria, for example, such as *Mycobacterium*, *Streptomyces*, and *Rhodococcus*, contain wax esters inside LDs, especially under stress or nutrition-depleted environment conditions (40, 41). It has been extensively documented that intracellular parasitic bacteria utilize LD for various reasons: obtaining nutrients or immune evasion (Nolan et al 2017). Our data showed that XH001 cells display enhanced saturated fatty acid production and formation of intracellular LDs when exposed to various stress conditions (**Figure 5**). Meanwhile, the increase in intracellular LDs is positively associated with XH001’s enhanced tolerance to starvation and oxidative stress (**Figure 6**). A recent study demonstrated that bacterial LDs bind to DNA via intermediary protein(s), thus protecting DNA and enhancing survival when exposed to genotoxic stressors (42). The improved resistance of XH001 to hydrogen peroxide (Figure 6), a genotoxic stressor, could potentially reflect the protective role of LDs in maintaining DNA stability, which is worth further examination. Thus, the enhanced accumulation of LDs could be a general response of XH001 to stress, which allows bacteria to better cope with adverse conditions for better survival. Our previous studies showed that the TM7x association reduced the XH001 growth rate and triggered the expression of stress-related genes (18). Therefore, the enhanced formation of LDs could be a strategic survival mechanism of XH001 in response to TM7x-induced stress.

Intriguingly, intracellular bacterial parasite infections often lead to the accumulation of host intracellular LDs, which are used by parasites for nutrients (Roingeard and Melo 2017). Thus, while the increased LD accumulation in XH001 when associated with TM7x may just be XH001’s general response to TM7x-induced stress, like in many other tested stress conditions, we cannot rule out the possibility that this could be a more specific parasite-bacterial host interaction. TM7x may induce LD production in XH001 to meet its growth demand. Genomic analysis reveals that *Saccharibacteria,* including TM7x, lack the capacity for *de novo* fatty acids synthesis (Castelle and Banfield 2018) while possessing membrane lipids rich in fatty acids (Probst et al 2020). Furthermore, studies have reported that intracellular parasites such as *Toxoplasma* can scavenge host lipid droplets for their intracellular development (Nolan et al 2017). Hence, we hypothesize that CPR bacteria might symbiotically obtain lipids or lipids building blocks from host organisms, an intriguing question that warrants further investigation.

Raman peaks at 1128 cm^-1^ and 1174 cm^-1^ further suggest that the major lipid molecules inside those lipid droplets might be saturated lipids such as polyhydroxybutyrate (Furukawa et al 2006), myristic acid, stearic acid (Czamara et al 2015). For future work, mass spectrometry analyses can be employed to determine which specific saturated lipids account for the major component of these LDs. We also found that co-culture XH001/TM7x has significantly higher cytochrome c content compared to host cells XH001 alone (**Figure 4E**). And the difference was not from the direct addition of TM7x as pure TM7x culture did not exhibit apparent Raman intensity from those peaks (**Supplementary Figure 2**). Cytochrome c, the ubiquitous heme-containing protein existing in all the life domains, is a key protein in the respiratory electron transfer chain of cells (Thöny-Meyer 1997). The higher content of cytochrome c suggests an enhanced respiration process for the host bacteria in the co-culture XH001/TM7x. Further studies regarding the correlation between enhanced respiratory activity with potential bacterial stress are needed to unravel this issue. Besides cytochrome c and saturated lipids, we also noticed other spectroscopic differences between XH001 cells and XH001/TM7x in amide I/III (proteins, 1660 cm^-1^/1250 cm^-1^). Raman intensities of amide I/III peaks in XH001 cells are higher than that of XH001/TM7x cells, implying that TM7x association elicits a metabolic activity shift inside the host cells.

In this study, through super-resolution fluorescence imaging and label-free Raman spectroscopy, we revealed some intriguing new features of XH001 host cells induced by *Saccharibacteria*, such as changes in cell membrane fluidity and the enhanced formation of LDs in XH001 as a result of symbiotic interaction with TM7x. However, due to the ultrasmall size of TM7x and the still limited lateral resolution of airyscan detector, future work is warranted to acquire nanoscale visualization of TM7x cell structures using advanced microscopic techniques such as stochastic optical reconstruction microscopy (19) or expansion microscopy (43). Another interesting direction to pursue is to find out the dominant lipid biomolecules that play an essential role in the attachment between TM7x and host cells. Taken together, our findings provide a new perspective to understand the evolutionary strategies of *Saccharibacteria* and its host bacteria.

## Materials and Methods

### Bacterial strains and growth conditions

XH001 monoculture and XH001/TM7x co-culture were isolated from the oral cavity as described in the previous published study (He et al 2015). Strains were cultured in brain heart infusion (BHI, Fisher Scientific, NH) at 37 °C under different oxygen conditions as specified in the main text: anaerobic (0 % O_2_, 10 % CO_2_, 5 % H_2_, balanced with N_2_), microaerophilic (2% O_2_, 5 % CO_2_, balanced with N_2_), and atmospheric conditions (∼21 % O_2_, 0.04 % CO_2_, 0.9 % Ar, 78 % N_2_). To acquire growth kinetics and phase contrast images, three independent cell cultures were grown under the specified oxygen condition for two passages (1 ml culture inoculated into 10 ml BHI and incubated 24 h each) before being reinoculated into 20 ml fresh BHI. The optical density at 600 nm (OD_600_) was measured using a spectrophotometer (ThermoFisherScientific, MA).

*Fusobacterium nucleatum* strains were cultured in Columbia broth (CB, Fisher Scientific, MA) at 37 °C under anaerobic condition (0 % O_2_, 10 % CO_2_, 5 % H_2_, balanced with N_2_) for two passages.

*Streptococcus mutans* UA159 and methicillin-resistant *Staphylococcus aureus* (MRSA) were cultured inside BHI broth at aerobic condition (∼21 % O_2_, 0.04 % CO_2_, 0.9 % Ar, 78 % N_2_) for two passages.

### Minimum inhibitory concentration of H_2_O_2_ against multiple bacteria

For XH001 cells, *Fusobacterium nucleatum* ATCC23726, *Fusobacterium nucleatum* ATCC10953, an initial bacterial inoculum with an OD_600_ of 0.1 was prepared and added to a sterile 96-well plate (Fisher Scientific, MA). 880 µM of H_2_O_2_ was added to the front row of a 96-well plate, then a two-fold serial dilution was performed with a resulting concentration of 880 µM, 440 µM, 220 µM…0 µM. After overnight incubation inside the microaerophilic chamber (XH001 cells) and anaerobic chamber (*Fusobacterium nucleatum* cells), the OD_600_ of the 96-well plate was recorded by a plate reader (iMark^TM^ microplate absorbance reader, Bio-Rad). The minimum inhibitory concentration (MIC) was determined as the lowest concentration at which no visible growth occurs. Sub-MIC bacterial inoculum was fixed with 10% formalin and washed three times with 1×PBS for the Raman spectra acquisition and fluorescence imaging.

For *Streptococcus mutans* and MRSA, all steps mirrored the aforementioned procedure, with the exception that the bacterial inoculum featured a concentration of 1×10^5^ cells/ml, the initial H_2_O_2_ concentration stood at 8.8 mM, and the cells were incubated within an aerobic chamber.

### Starvation assay

XH001 cells were initially cultured in a microaerophilic chamber for two passages. Subsequently, two distinct groups were prepared: an untreated group and a prestress group. The cells in each group were diluted and cultured overnight, with the untreated group incubated inside a microaerophilic chamber and the prestress group within an anaerobic chamber. Following this, the cells were harvested, subjected to two rounds of washing with sterile 1×PBS, and ultimately suspended in PBS within the microaerophilic chamber. Starvation happened over time, samples were collected at each specific point, and subjected to the following colony-forming unit (CFU) assay, susceptibility test towards H_2_O_2,_ and Raman spectra acquisition.

### Fluorescence microscopy

For BODIPY C_1_, C_12_ fluorescence imaging, cells were fixed, washed with 1×PBS, and stained with BODIPY C_1_, C_12_ (Thermo Fisher Scientific, MA) with a working concentration of 1.5 µM for 30 min. After that, cells were washed twice with 1×PBS, and sandwiched below a cover glass (Fisher Scientific) and glass slide (Fisher Scientific, NH). Fluorescence images were acquired at an excitation wavelength of 488 nm and an emission window from 500-550 nm. Confocal laser scanning microscopy was conducted by a ZEISS LSM880 (Carl Zeiss AG, Germany) system with a 63× (NA=1.4) oil immersion objective.

For LipidSpot fluorescence imaging, live cells (XH001 cells, XH001/TM7x cells) were stained with LipidSpot (Biotium) at a 1×working concentration for 3-5 hours. After that, cells were immediately collected to acquire the fluorescence images at an excitation wavelength of 488 nm and an emission window from 500-550 nm under a ZEISS LSM880 confocal laser scanning microscope.

For Laurdan-based membrane fluidity measurements, Laurdan (6-Dodecanoyl-2 Dimethylaminonaphthalene Thermo Fisher Scientific, MA) was dissolved in DMF (Sigma Aldrich, MA) and a final concentration of 1% DMF was maintained in the medium for better solubility. Cells were washed two times with prewarmed PBS and were further incubated with Laurdan for 30 min. After that, cells were washed two times with prewarmed PBS and sandwiched between a cover glass and a cover slide. Laurdan fluorescence intensities were measured at 460±5 nm and 500±5 nm upon excitation at 405 nm under a ZEISS LSM880 confocal laser scanning microscope with the airyscan detector. Images were acquired with a 63× (NA=1.4) oil immersion objective. Laurdan generalized polarization (GP) value was calculated using the formula GP = (I_460_-I_500_)/(I_460_+I_500_).

Airyscan imaging was performed with a confocal laser scanning microscope ZEISS LSM 880 equipped with an Airyscan detection unit. To maximize the resolution enhancement, we used high NA oil immersion Plan-Apochromat 63× (NA=1.40) Oil Corr M27 objectives (Zeiss, Germany). All imaging was performed using Immersol 518 F immersion media (n_e_ = 1.518 (23 °C); Carl Zeiss). Detector gain and pixel dwell times were adjusted for each dataset keeping them at their lowest values in order to avoid saturation and bleaching effects.

For Airyscan processing (super-resolution fluorescence imaging), Zen Black software was used to process the acquired images. The software processes each of the 32 Airy detector channels separately by performing filtering, deconvolution, and pixel reassignment in order to obtain images with enhanced spatial resolution and improved signal-to-noise ratio. This processing includes a Wiener filter deconvolution with options of either a 2D or a 3D reconstruction algorithm (details are described elsewhere (Huff 2015)). In our analysis, we applied the Airyscan Processing Baseline Shift and further used the 2D or 3D reconstruction algorithm either at the default filter setting or at filter settings as noted.

All acquired images were analyzed through FiJi (NIH) and CellProfiler.

### Measurement and analysis of Raman spectra

Cultured cells were fixed in 10% formalin (neutral buffered, Sigma Aldrich, MA) for 60 min and were washed three times with sterile water. Before Raman measurements, the fixed cells were dropped onto an aluminum-coated Raman substrate to be air-dried. Raman spectra were acquired using an HR Evolution confocal Raman microscope (Horiba Jobin-Yvon, France) equipped with a 532-nm neodymium-yttrium aluminum garnet laser. The laser power on cells was 8 mW after attenuation by neutral density filters (25%). An objective with a magnification of 100× (NA=0.6) was used to focus single cells with a laser spot size of ∼1 μm^2^, and Raman scattering was detected by a charge-coupled device cooled at −70 °C. The spectra were acquired in the range of 300 cm^−1^ to 1800 cm^−1^ with 600 grooves per mm diffraction grating. The acquisition parameters were 10 s with 2 accumulation per spectrum, at least 9 spectra from three biological replicates. All obtained Raman spectra were preprocessed by comic ray correction and polyline baseline fitting with LabSpec 6 (Horiba Scientific, USA). Spectral normalization was done by vector normalization of the entire spectral region. The selection of vector normalization was undertaken with the aim of rectifying general instrumentation fluctuations, as well as mitigating the influence of sample and experimental variables (such as the thickness of the sample) while minimizing interference with the inherent characteristics of the biological content. Data analysis, statistics, and visualization were performed by OriginLab (OriginLab Corporation, MA) and RStudio 2023.03.0+386 environment using in-house scripts.

### Statistical analysis

Two-tailed Student’s unpaired *t*-test and one-way analysis of variance (ANOVA) were used to determine whether there is any statistically significant difference between groups (\**p* < 0.05, \*\**p* < 0.01, \*\*\**p* < 0.001, \*\*\*\**p* < 0.0001).

## Acknowledgements

**Funding**

This work was supported by R01DE023810 and R01DE030943 to X.H. and, in part, by T90DE026110 to P.-T.D.

**Author contributions**

P.-T.D. conceived the research and made the accidental discovery of LDs formation inside XH001 cells. P.-T.D., J.T., and L.C. prepared the samples. P.-T.D. performed experiments, data acquisition, image acquisition, and data analysis. KJ.K.K. provided suggestions on the data analysis part of Raman spectra. P.-T.D., W. S., and X. H. co-wrote the manuscript. X. H., W. S., B.B., and J. S. M. contributed to the constructive suggestions over the experiments. All authors discussed the results and contributed to the writing of the manuscript.

**Competing interests**

The authors declare that they have no competing interests.

**Data and materials availability**

All data needed to evaluate the conclusions in the paper are present in the paper and/or the Supplementary Materials. Additional data related to this paper may be requested from the authors.

## Supplementary Figures

**Supplementary Figure 1.**
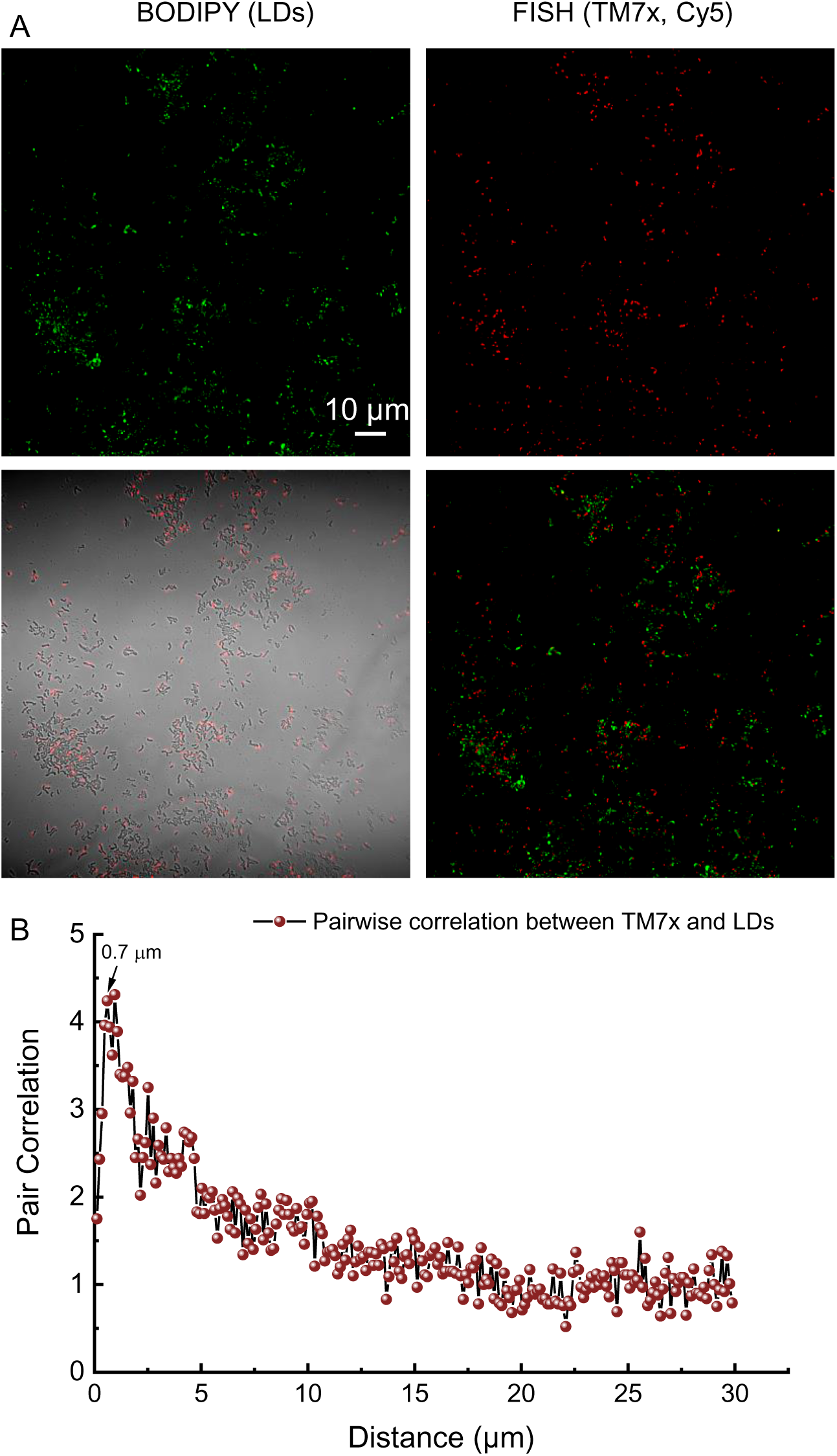
Spatial arrangement between lipid droplets and TM7x (FISH staining) in the co-culture between XH001 and TM7x. **A.** Confocal fluorescence images of BODIPY-labeled XH001/TM7x cells and TM7x (after fluorescence in situ hybridization). Scalar bar = 10 µm. **B.** Pairwise correlation from the above two channels. The highest pair correlation value was labeled at a dipole distance of 0.7 µm.

**Supplementary Figure 2.**
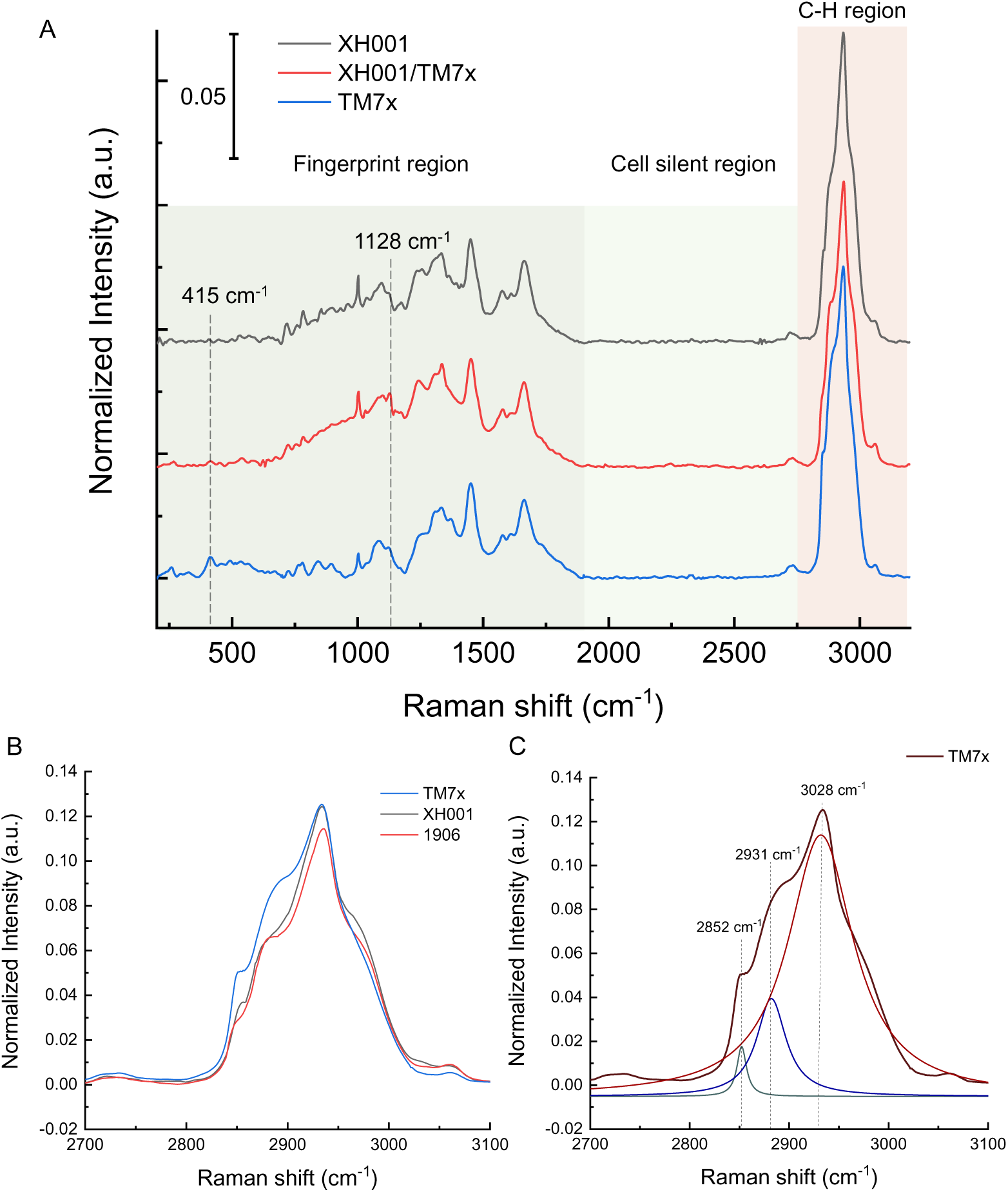
Averaged Raman spectra of XH001, co-culture XH001/TM7x with prophage xhp1, and TM7x. **A**. Averaged Raman spectra of XH001, XH001/TM7x, and TM7x cells in the range of 200 to 3200 cm^-1^. Three different regions were highlighted with different colors (fingerprint region, cell silent region, and C-H region). B. Comparison of the Raman spectra in the C-H region from the above three cells. **C**. Lorentzian fitting of Raman spectrum in the C-H region of TM7x cells. Three major components were decomposed under a Lorentzian fitting.

**Supplementary Figure 3.**
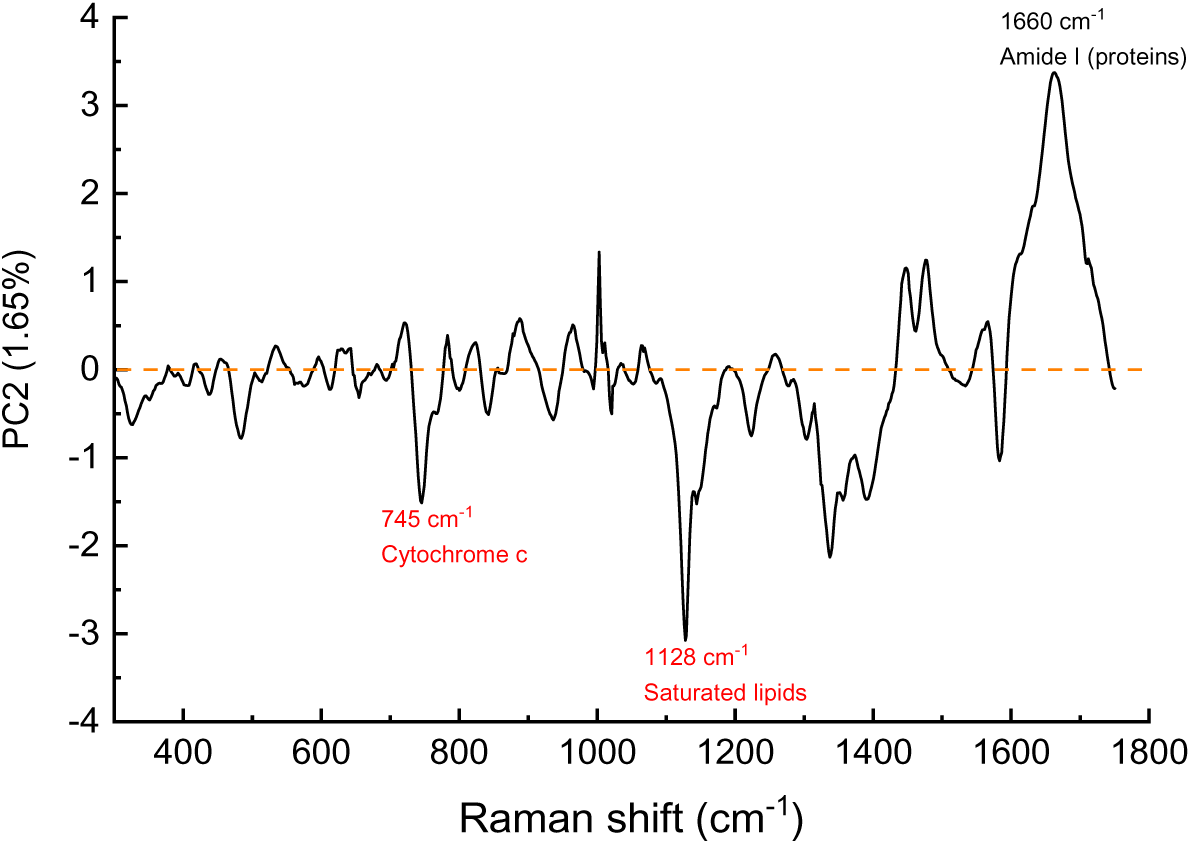
Spectral information of principal component 2 which contributes to 1.65% of the total variation. Peaks of interest were highlighted in the plot.

**Supplementary Figure 4.**
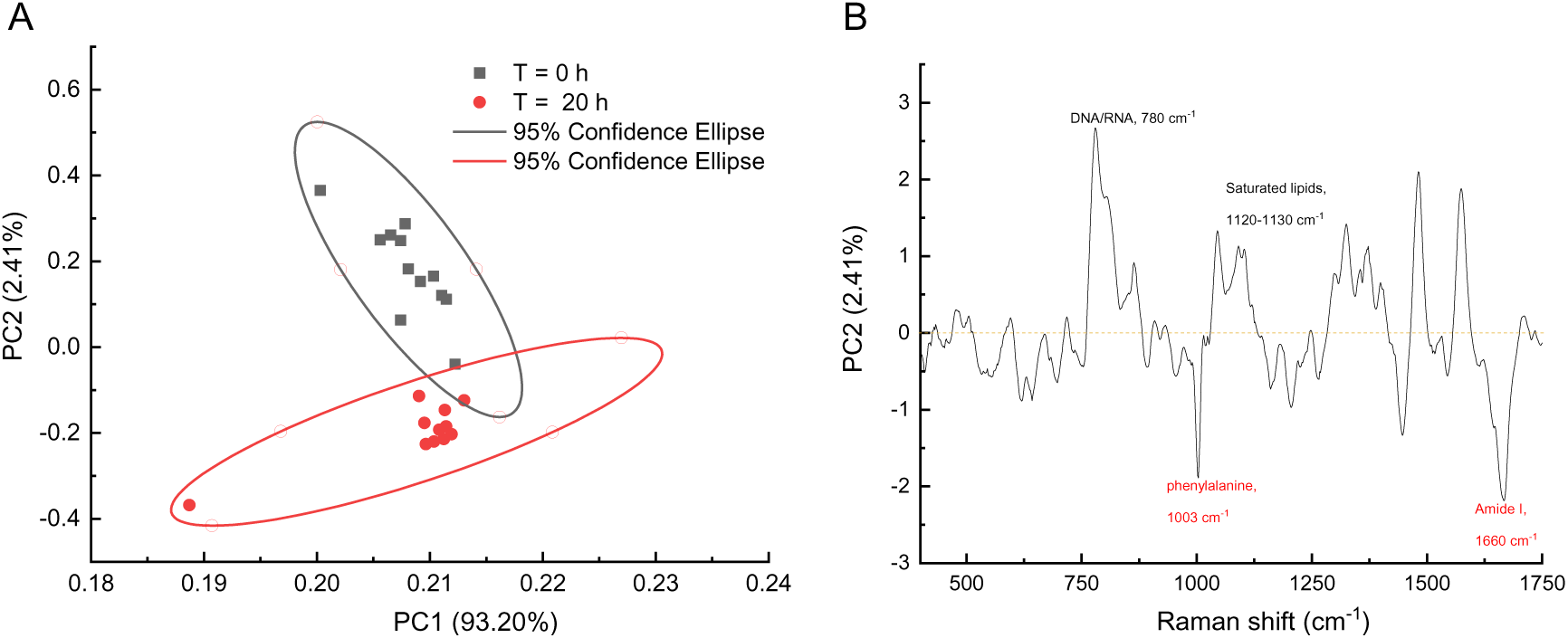
Principal component analysis of Raman spectra acquired from pre-stressed XH001 cells at different starvation times. Plots were depicted from principal component 1 (93.20% of the total variation) and 2 (2.41% of the total variation). 95% confidence intervals were represented by ellipses. Peaks of interest were highlighted in the plot.

**Supplementary Figure 5.**
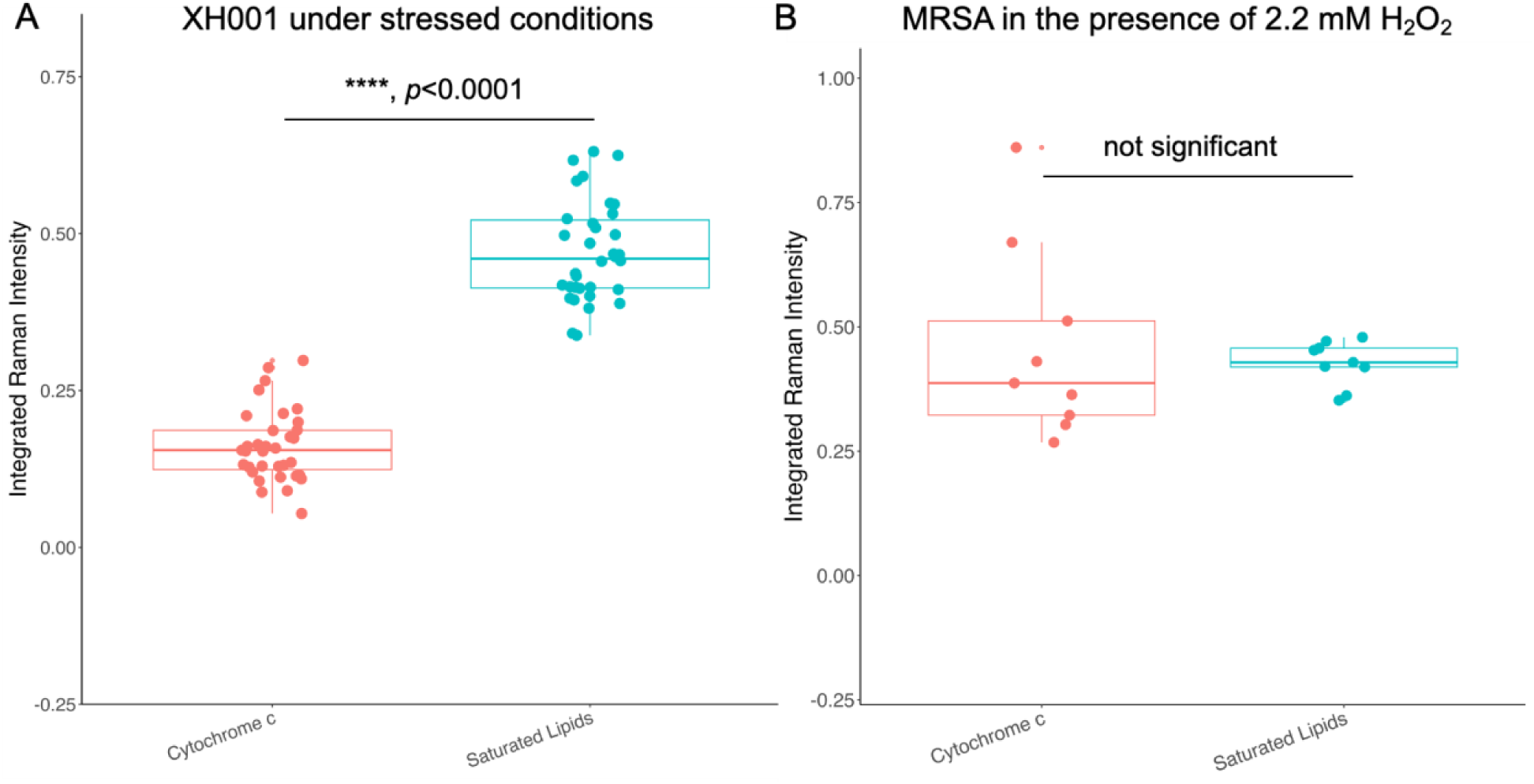
Quantification of the integrated Raman Intensity from cytochrome c and saturated lipids in the case of XH001 cells and *Staphylococcus aureus* cells under stressed conditions. Each dot is from a single Raman spectrum. Statistical analysis was achieved by a two-tailed student unpaired *t*-test.

